# Hyaluronan Hydrogel “Safety Nets” for 3D Cell Culture Applications

**DOI:** 10.1101/2023.08.12.553089

**Authors:** Russell T. Wilson, Andrei Bonteanu, Daniel A. Harrington

## Abstract

Polymeric hydrogels can mimic features of native extracellular matrix (ECM), and their facile production and characterization enable customizable cell encapsulation and 3-dimensional (3D) culture. These have applications relevant to dentistry such as soft tissue engineering, cancer modeling, and drug screening. Hyaluronan (HA), a biologically-derived polymer found in native ECM, offers an optimal platform for this use, as it can be modified covalently with adhesive ligands, enzyme-degradable crosslinkers, and other biorelevant moieties, yielding tailored physical properties and biological response. Cell-directed degradation of these encapsulating matrices can be a prerequisite for phenotype preservation, but degradation kinetics may not match desired timelines for drug screening applications. This study focused on optimizing hydrogel composition, network structure, and gelation kinetics to preserve z-distribution of physiologically relevant cells within high-throughput microfluidic plates. Hydrogels were formed from aqueous solutions of thiolated HA (HA-SH), bifunctional acrylated poly(ethylene glycol)-peptide crosslinkers, and pendant acrylated peptides (RGD or YIGSR sequences) to support cell adhesion. By varying the absolute crosslinker concentration and relative proportions of high:low crosslinker degradation kinetics, hydrogels could be tuned to desired moduli (G’: ∼10-120 Pa) and enzymatic degradation rate. Ratios of high:low degradable crosslinkers, at equivalent total crosslinker concentration, minimally impacted final modulus or gelation rate. Similarly, pendant adhesive ligands were swapped easily with negligible impact on hydrogel physical properties. Hydrogels with a 50:50 ratio of high:low degradable crosslinkers provided a “safety net” that preserved z-distribution of encapsulated bone marrow-derived fibroblasts, while maintaining expected phenotype. A comparable system supported the 3D co-culture of primary human salivary epithelial and mesenchymal cells within a perfusable microfluidic multiwell plate. This customizable bottom-up construction reduced confounding factors encountered in hybridoma-derived protein matrices, enabling modular customization of physiologically relevant, yet reproducible matrices to replicate native ECM. Our optimized model demonstrates workflows to improve future pharmaceutical screens, and tailor tissue engineering applications.

## Introduction

Three-dimensional (3D) cell culture has been used widely over decades for multiple research applications including cancer modeling, drug screening, trauma repair, and tissue engineering (Gomez-Florit et al. 2020; Chyzy and Plonska-Brzezinska 2020; Boucherit, Gorvel, and Olive 2020; Fong, Harrington, et al. 2016; Highley, Prestwich, and Burdick 2016). The definition of 3D culture is subject to broad interpretation, and variations often are identified by the amount, type, and characteristics of a supportive extracellular matrix (ECM). In the present work, we constructed semi-synthetic hydrogel matrices that mimic the mechanical and biological cues of native soft ECM and enable encapsulated cells to be suspended in 3D space. Such systems enable phenotypic cell-ECM interactions, such as cell migration, matrix degradation, nutrient diffusion, and cell receptor interaction with matrix ligands, among many others(Baruffaldi et al. 2021). Multiple cell types can be encapsulated within a single culture and assume complex morphologies, giving insight into how they interact with and signal each other in both healthy and diseased states (Sablatura et al. 2023; Boucherit, Gorvel, and Olive 2020; Langhans 2018; Fong, Harrington, et al. 2016).

The options available for ECM mimics vary broadly, across synthetic and naturally-derived options, and researchers’ choices rely heavily on their potential application and on any necessary characteristics of the matrix. Naturally-derived options, including basement membrane extracts and collagen I, offer expansive biological content that cells recognize, but permit minimal compositional control and may suffer from issues of reproducibility or inconsistency in structural properties (Aisenbrey and Murphy 2020; Antoine, Vlachos, and Rylander 2014; Liu et al. 2011; Peterson 2008). Our contrasting approach leverages hyaluronan (HA) polymer, which is present throughout native ECM, and offers a biologically-relevant foundation on which to build customized features (Appendix Figure 1). Thiolated HA (HA-SH) serves as a modular backbone that can be combined with thiol-reactive acrylated crosslinkers to form stable hydrogels and decorated with pendant acrylated epitopes to improve cell adhesion and influence other behaviors (Highley, Prestwich, and Burdick 2016; Misra et al. 2015; Xu et al. 2014; Burdick and Prestwich 2011; Shu et al. 2004; Miranti and Brugge 2002).

Matrix metalloproteinase (MMP)-labile peptides, with short poly(ethylene glycol) (PEG)-acrylate functionality on each side, have been employed as bifunctional crosslinker molecules to enable initial network formation and facilitate later cell-directed remodeling (Lutolf et al. 2003). This dynamic interaction with the matrix can be carried out by a wide range of cell types, as this degradable crosslinker is derived from a mutated component of type I human collagen, and has reliable degradation by multiple MMPs (Lutolf et al. 2003). However, some cell types or conditions may exhibit overly aggressive degradation of such matrices, leading to early cell “sinking”, in a manner that is premature for evaluation timepoints in an *in vitro* assay. A tailored hydrogel, with well-defined mechanical properties and tempered enzymatic lability, could preserve metered cell motility and stretching, while avoiding a rapid collapse of cells to the matrix base. A controlled ratio of crosslinkers with variable degradation profiles may act as a “safety net” (Figure 1), which will prevent excessive degradation and subsequent softening of the hydrogel.

**Figure 1:**
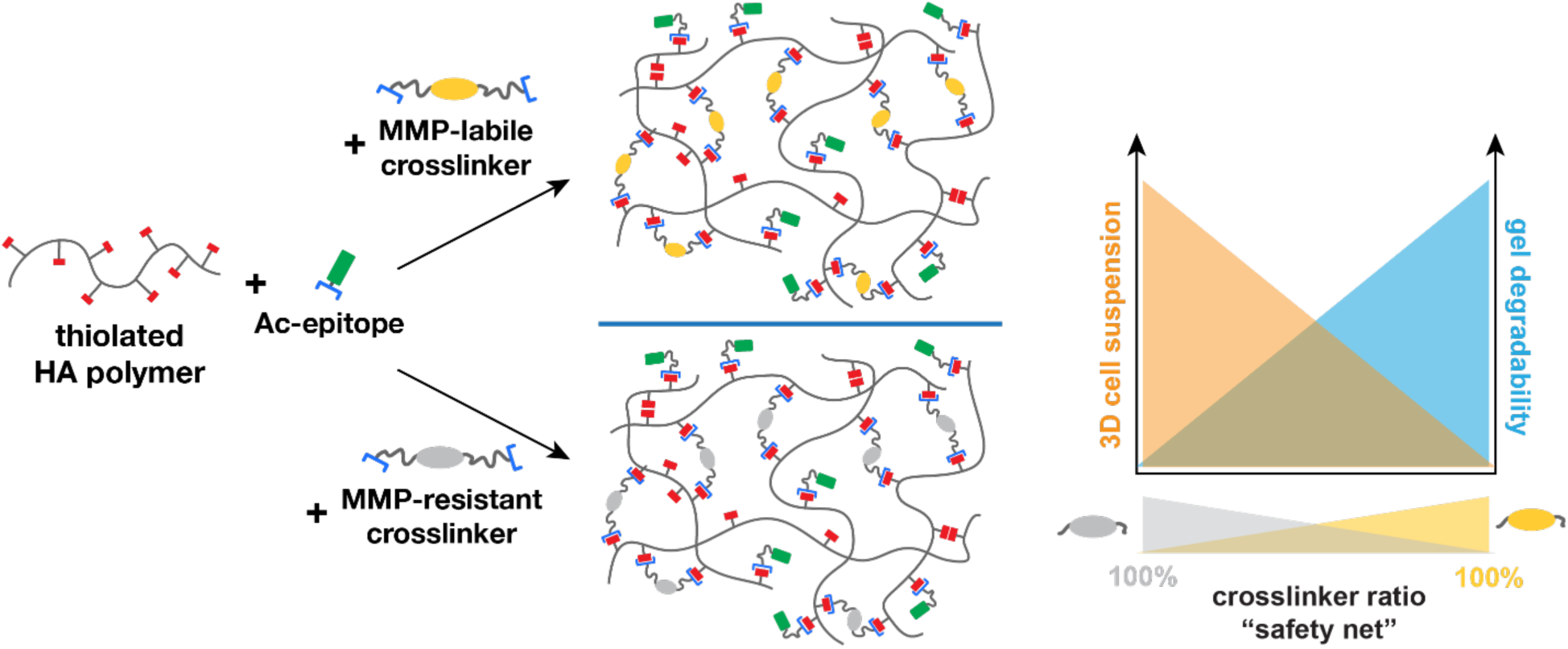
HA-SH hydrogels can be tuned with either matrix metalloproteinase (MMP) labile crosslinkers, MMP resistant crosslinker, or a combination of both. This work aimed to identify a ratio of high:low MMP degradable crosslinkers that retains hydrogel degradability that some cells (e.g. fibroblasts) require in order to preserve phenotypic stretching and migration, while avoiding complete dissolution of the in vitro matrix. An optimized ratio of these crosslinkers could preserve cells in 3D suspension while retaining their phenotype, thereby forming a “safety net” in the hydrogel.

In this work we assessed via rheology the gelation properties of modular HA-SH hydrogels by exchanging crosslinkers with varying degradation profiles, at varying densities, to determine their later applicability toward 3D culture and drug screening platforms. Similarly, pendant biorelevant epitopes were exchanged into the hydrogel network to assess the modularity of the system. Ratios of crosslinkers with similar molecular structure but varying fast/slow MMP degradation profiles were incorporated at equivalent total crosslink density to assess changes in initial hydrogel properties and long-term resistance to MMP degradation. Lastly, we tested the application of this model toward a microfluidic drug screening platform, with co-cultures of multiple salivary cell types, to demonstrate viability of the modular hydrogel model. The modifications introduced in this work represent a critical need toward optimization of HA-SH hydrogels for cancer modeling in high-throughput drug screening. Moreover, these protocols and workflows for optimizing hydrogel physical properties will be applicable to other 3D cell culture applications, ranging from tumor modeling and drug screening to tissue engineering and trauma repair.

## MATERIALS AND METHODS

Details of materials and methods are provided in the Supplemental Appendix. Briefly: HA-SH and all peptides were commercially sourced. Pendant adhesive peptides RGD and YIGSR were synthesized with acryl groups at the N-terminus. Crosslinkers were generated from “PQ” and “PQ*” peptides, (PQ: KGGGPQG↓IWGQGK; PQ*: KGGG<u>D</u>QGI<u>A</u>G<u>F</u>GK, in which ↓ represents the cleavage site), by reacting each with amine-reactive SVA-PEG-Ac. This yielded bi-acrylated, bi-PEGylated PQ and PQ* crosslinkers, with high or low susceptibility to MMP degradation (respectively), yet with very similar molecular weights for the final bi-PEGylated products. A diagram of these components is shown in Appendix Figure 1.

## RESULTS

### HA-SH Crosslinking Kinetics Increases with pH

Hydrogel selection for 3D cell culture is informed by multiple factors, including the kinetics of gelation, the ability to encapsulate cells easily with high viability, the range of accessible mechanical properties (initial, and over time), and the replication of biological cues found in the native tissue. To set a foundation for understanding the base gelation kinetics of the HA-SH system, we assessed the impact of pH on both the rate of gelation, and the ultimate G’ value. These HA-SH systems react with poly(ethylene glycol)diacrylate (PEGDA) oligomers via a Michael addition to form crosslinked hydrogel networks(Ghosh et al. 2005; Shu et al. 2004). Prior work demonstrated the pH sensitivity of this thiol-acrylate reaction (Lutolf et al. 2001). Given our target use of cell encapsulation, such pH ranges must support high cell viability, thus restricting our exploration to values between pH 7.0 and 8.0.

A range of crosslinkers and pendant epitopes (Appendix Figure 1), at multiple concentrations, were reviewed throughout this study. To assess the impact of pH on all of these conditions, we established a representative hydrogel network condition using an intermediate concentration of PQ crosslinker (0.55 mM) and 3.0 mM of pendant Ac-RGD. The pH of each hydrogel precursor solution was preset, between 7.0-8.0, immediately prior to their combination, loading onto the rheometer, and analysis over time, up to 60 minutes. As shown in Appendix Figure 2, all conditions formed a gel, and gelation rate increased substantially with pH. All conditions exhibited a lag time, prior to a linear region of G’ increase, and lag time increased as pH decreased. Only the pH 8.0 condition exhibited a near-sigmoidal response over time, with accelerated crosslinking within 10 minutes of mixing, and a near-plateau modulus by 60 minutes. Because these kinetics are optimal for rapid solution dispensing and maintenance of suspended cells, pH 8.0 was used for all subsequent studies on hydrogel network structure.

### Hydrogel Storage Modulus (G’) Increases with Crosslinker Concentration

We next sought to determine how storage modulus (G’) and gelation kinetics scaled with crosslinker concentration. At pH 8.0 and a baseline level of 3.0 mM Ac-RGD, we varied the concentration of PQ crosslinker from 0.3-0.6 mM. As anticipated, increasing crosslinker concentration led to a higher final G’ value, and increased gelation rate, yet the times to initiate and complete gelation were similar across all concentrations tested (Figure 2A).

**Figure 2.**
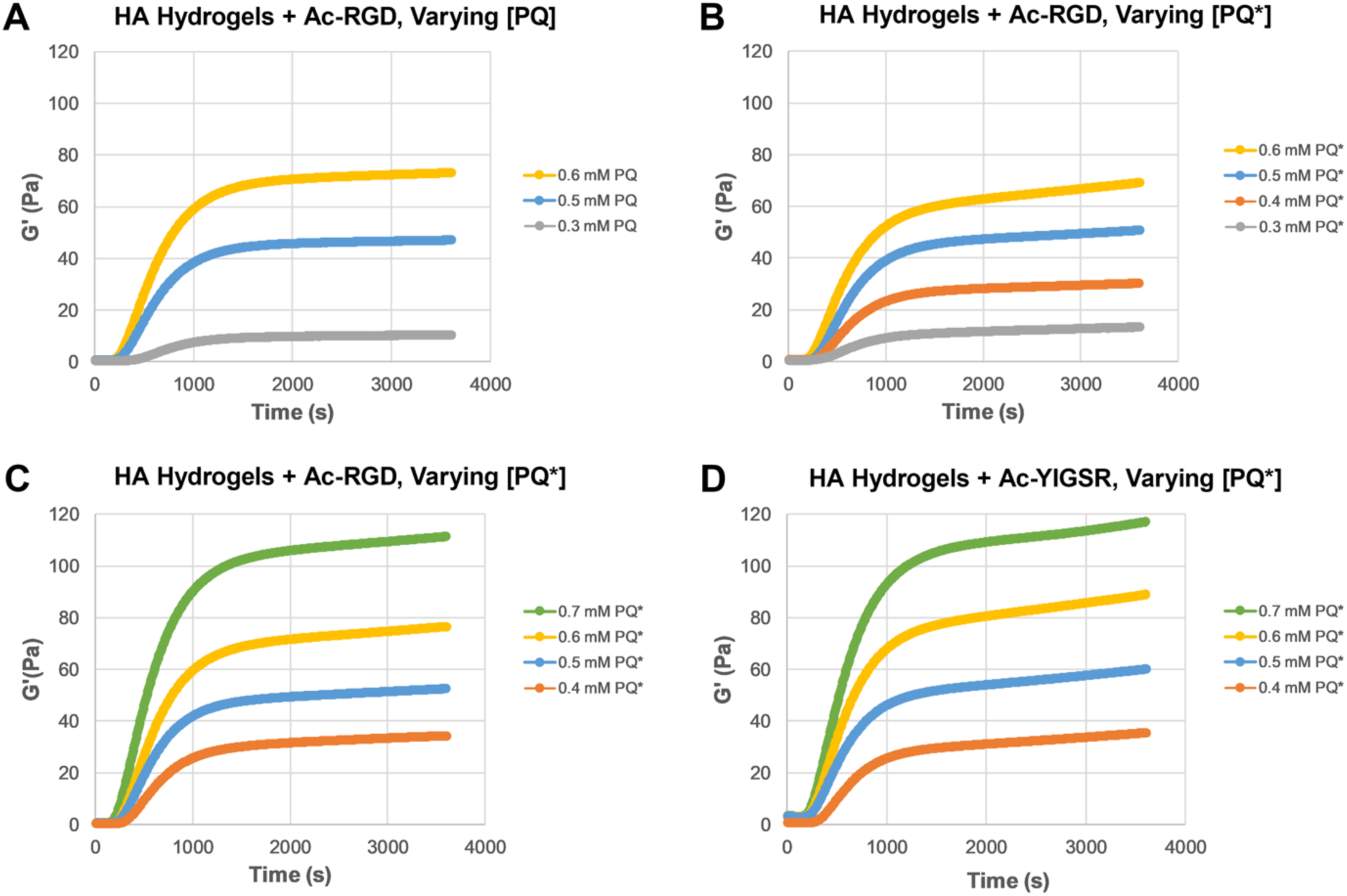
Both PQ and PQ* crosslinkers, and Ac-RGD and Ac-YIGSR pendant epitopes, are readily exchanged in HA hydrogels to yield similar mechanical properties and kinetics, scaling with crosslinker concentration. Rheologic assessment of HA systems, immediately after crosslinker addition, is shown. (A) PQ and (B) PQ* crosslinkers were added to HA-SH, with 3.0 mM of Ac-RGD pendant epitope, at the crosslinker concentrations noted in the legends. HA-SH systems containing 3.0 mM of (C) Ac-RGD or (D) Ac-YIGSR pendant epitopes were crosslinked using multiple concentrations of PQ* noted in each legend. Within modest experimental variation, all systems behaved similarly, with similar rates of gelation and plateau levels.

### Exchanging PQ for PQ* Crosslinker Yields Equivalent G’ Moduli

The PQ and PQ* crosslinkers have very similar chemical structures, except for small exchanges in the central peptide sequences that define susceptibility to MMP degradation. We anticipated that these crosslinkers could be swapped evenly at the same molar concentration, and generate minimal impact on hydrogel gelation rate or final modulus. To test this hypothesis, the conditions of the previous experiment were replicated exactly, exchanging PQ crosslinkers with PQ* variants, at equivalent molar concentrations. As shown in Figure 2B, the resultant G’ moduli for the PQ* system were consistent with the values for equivalent concentrations with PQ. These data provide a basis for later ratios of the two crosslinkers to generate “safety net” networks with tunable lability to MMPs.

### Exchanging Pendant Peptide Epitopes Maintains Gelation Behaviors

In addition to modular exchange of crosslinkers, the HA-SH hydrogel system offers a modular exchange of pendant epitopes, with a maximum value defined by the thiolation level of the HA-SH backbone. Customization of these hydrogels may require integration of different, or multiple, epitopes. We hypothesized that small, similarly-sized epitopes, such as the integrin ligands RGD and YIGSR, could be swapped into the network with minimal effect on mechanical properties or gelation. Using the PQ* crosslinker, at equivalent concentration ranges from 0.4-0.7 mM, hydrogels were prepared with either 3.0 mM Ac-RGD (Figure 2C) or equivalent molar concentrations of Ac-YIGSR (Figure 2D). Hydrogels with Ac-YIGSR showed similar, albeit slightly higher, G’ values than hydrogels with Ac-RGD.

### “Safety Net” Ratios of Crosslinkers Yield Similar Hydrogel Moduli

We next sought to explore the effects of a “safety net” architecture on hydrogel properties, using both PQ and PQ* crosslinker at varying ratios. Hydrogels were prepared with total crosslinker concentration ([PQ + PQ*]) at 0.4 mM (Figure 3A), and repeated at 0.5 mM (Figure 3B). Ac-RGD was incorporated at the standard 3.0 mM. Both experiments showed that PQ:PQ* ratios of 0, 25, 50, and 100% PQ* yielded similar gelation behavior at both crosslink densities, yielding similar initial lag times, gelation rates, and final G’ moduli within a range of ∼ 10 Pa, with higher G’ on the 100% PQ* sample.

**Figure 3.**
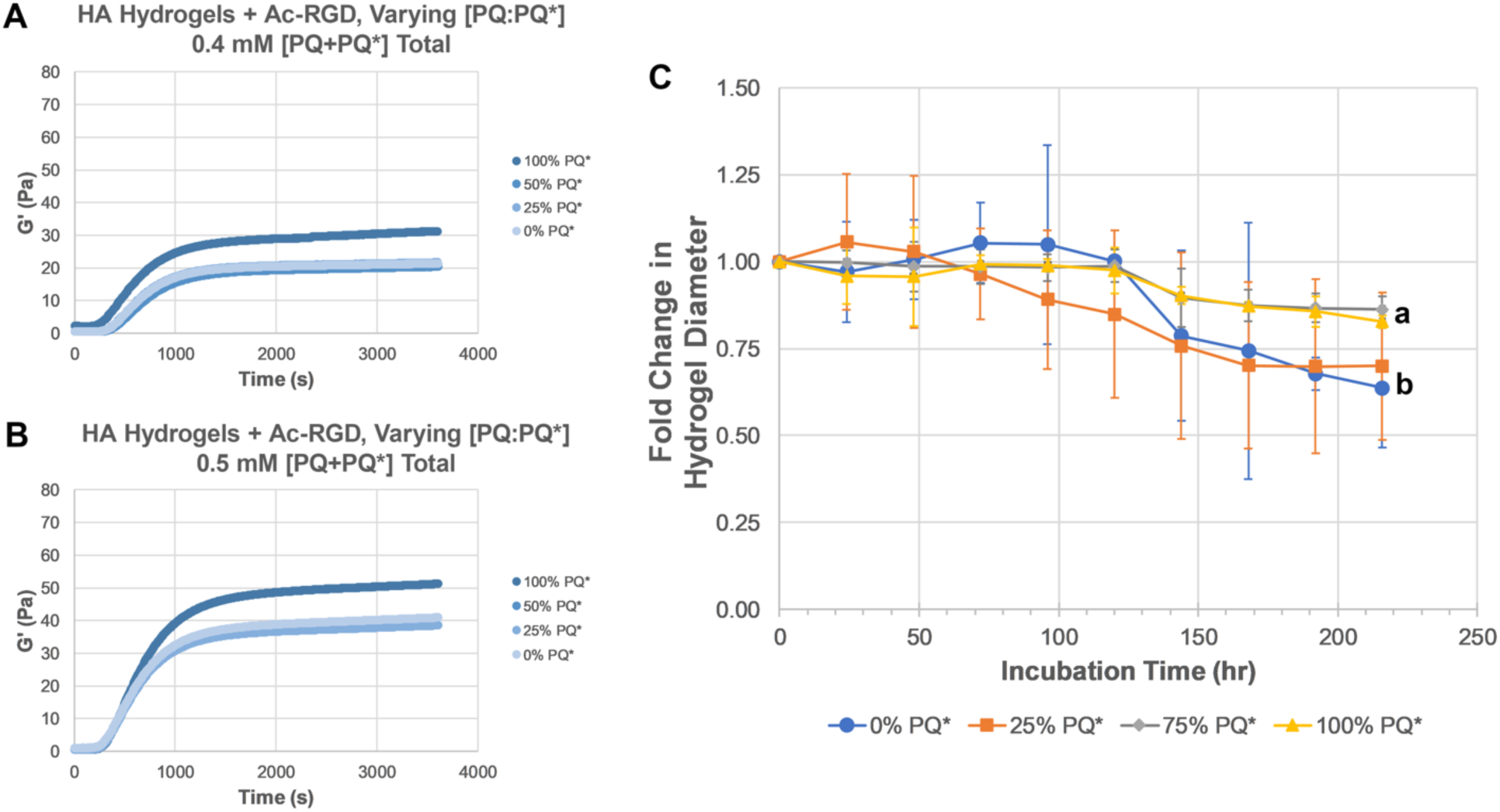
“Safety net” hydrogels, using ratios of PQ and PQ* crosslinkers, yield similar mechanical properties and improve resistance to MMP-based degradation. Hydrogels were prepared at multiple PQ:PQ* ratios, held at total crosslink densities of (A) 0.4 mM and (B) 0.5 mM. Ratioed variants within each crosslink density maintained similar kinetics and final moduli, with a slight increase in the 100% PQ* variants. (C) Hydrogel discs for each ratio were cast in silicone molds and incubated to 9 days, submerged in 1 µg/mL human MMP-1 solution. Changes in the hydrogel diameters, normalized to the starting diameter, were recorded from daily images. N=3 repeats, error bars: ± 95% confidence interval. 1-way ANOVA comparison with Tukey’s multiple comparisons at the final timepoint (216 hr) showed statistically significant difference between means (p<0.05) for compositions marked with different letters (a,b).

### “Safety Net” Hydrogels Resist MMP Degradation in Proportion to their PQ:PQ* Ratio

Hydrogel pucks were created using varying “safety net” ratios of PQ and PQ*, with total crosslinker concentration at 0.7 mM, and with 3.0 mM Ac-RGD. These pucks were submerged in human MMP-1 solution and diameter changes were measured over 9 days. Results (Figure 3C) showed that hydrogels with greater PQ* ratio (75 and 100% PQ*) exhibited less change in diameter when compared to hydrogels containing less PQ* (0 or 25% PQ*). 1-way ANOVA assessment at the final timepoint (t=216 hr) confirmed this difference, and a post-hoc Tukey’s test showed statistically significant differences (p<0.05) between the groups labeled as “a” (75% and 100% PQ*) and “b” (0% and 25% PQ*) on the graph in Figure 3C. Details of this comparison are provided in Supplemental Appendix Table 1. Additional analysis of these data using repeated measures (linear mixed effects) modeling further confirmed the combined impacts of hydrogel composition and time on changes in puck diameter.

### “Safety Net” Hydrogel Compositions Preserve Morphology and Z-Distribution of Fibroblasts

To test the application of these hydrogel variants with cells, we used HS27a bone marrow-derived fibroblasts, which have been shown previously to thrive only in PQ-functionalized matrices (Fong, Wan, et al. 2016). We hypothesized that matrix degradation would be tempered by substituting stoichiometric ratios of PQ crosslinker with a mutated PQ crosslinker sequence (PQ*) that exhibits less susceptibility to MMP-based degradation. Two HA hydrogel formulas with 25% and 50% PQ* incorporation (i.e. 25%:75% and 50%:50% ratios of PQ*:PQ) and 3.0 mM pendant RGD were used to encapsulate 5000 fibroblasts/μL within the gel channel (GC) of a perfusable microfluidic multiwell plate (Fig 4A). Two primary outputs were used in this assessment: fibroblast stretch ratio as an indication of phenotypic morphology, and intrusion of a high MW FITC-dextran solution from the perfusion channel (PC) into the GC, as a surrogate measure of hydrogel degradation. The latter method was assessed by recording a Z-stack of the FITC signal in the GC, reconstructing a 3D volume in Imaris software, and calculating a % intrusion (v/v), based on total volume available in the GC.

**Figure 4.**
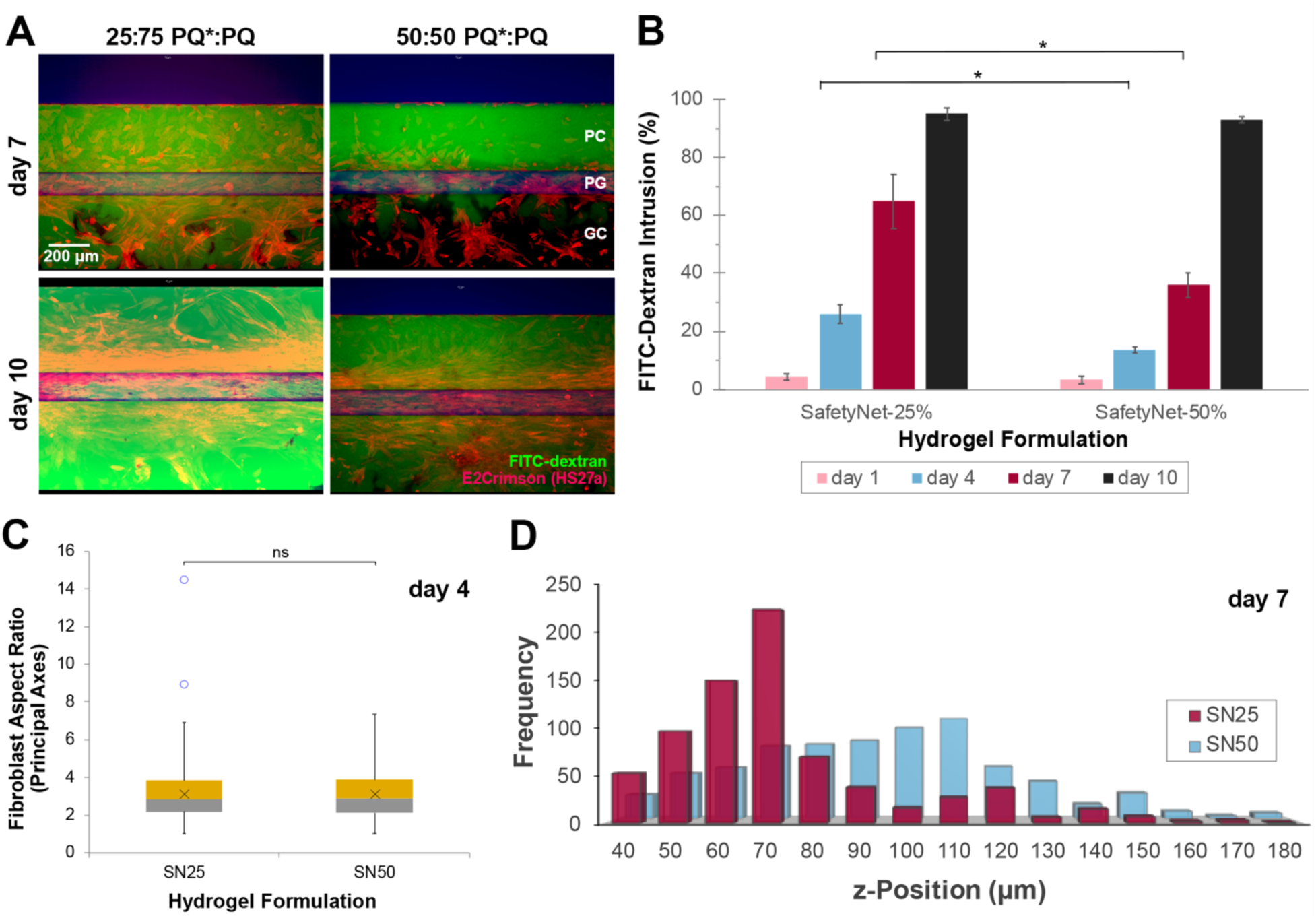
HA “safety net” crosslinker ligands delay fibroblast invasion and hydrogel degradation. (A) HS27a bone marrow-derived fibroblasts (constitutively expressing E2-Crimson, red) were encapsulated within hydrogels crosslinked with 25% PQ* or 50% PQ*, onto a microfluidic culture plate within the gel channel (GC). Cells exhibited phenotypic stretching within the gels. By 7 days of culture, a solution of 2MDa FITC-dextran (green), introduced into the adjacent perfusion channel (PC), diffused readily into the GC of the 25:75 hydrogel, yet was still restrained by the 50:50 hydrogel. Volume-based quantification of this FITC-dextran intrusion in (B) demonstrated differential response between the two materials at days 4 and 7. (C) Comparative analysis of fibroblast morphology at day 4 shows equivalent cell stretching in both matrices at this early timepoint. (D) A histogram of cell location in Z within the GC at day 7 identified that cells within SN25 hydrogel had preferentially fallen to a lower z position, while cells in SN50 maintained a more even distribution, centered toward the half-maximum (100 µm) of the 200 µm channel. * = p < 0.05. Error bars in (B) indicate standard deviation. Box plots in (C) show middle quartiles; x indicates average, whiskers indicate range, o denotes outliers beyond 2.2 standard deviations, ns = no statistical difference.

For both hydrogel compositions, the HS27a cells, constitutively expressing E2Crimson fluorescent protein for ease of imaging, demonstrated equivalent aspect ratios at day 1 (1.7, not shown) and day 4 (3.1, Fig. 4C). A fibroblast aspect ratio ≥ 3 was considered to reflect a stretched phenotype, and was observed from day 4 through day 10. For the 50:50 hydrogel, significantly less FITC-dextran intrusion was observed on both days 4 and 7 (14% and 36%, respectively) compared to the 25:75 hydrogel at the same timepoints (25% and 65%, respectively) (Fig. 4A, B). Additionally, in 50:50 hydrogels, fibroblast centroid positions on days 7 and 10 were found to be more evenly distributed in the z-dimension, and with a higher mean z-position, for cells in the 50:50 hydrogel safety net (Fig. 4D, Appendix Figure 3). Notably, this finding occurred even as the 50:50 hydrogel progressively degraded (observed as a gradual increase of FITC-dextran assay intrusion into the hydrogel).

### HA Hydrogels Support Co-Cultures of Salivary Epithelial Clusters & Fibroblasts in *In Vitro* Screening Plates

In an application of these hydrogel results toward an alternate use with co-cultured salivary epithelia and fibroblasts, human salivary epithelial cells (hS/PCs) were formed as clusters on a non-adhesive multiwell plate (Figure 5A) then seeded with a peripheral shell of human salivary fibroblasts (hSFs), which attached to and surrounded the epithelial cores. These clusters were suspended in HA-SH hydrogel precursor solution, with 2.0 mM Ac-RGD, 2.0 mM Ac-YIGSR, and 0.5 mM PQ crosslinker, and loaded onto a perfused microfluidic culture plate (Figure 5B). After 7 days in culture, the salivary co-cultures remained highly viable within the hydrogel matrices, and the peripheral cells extended into the surrounding hydrogel (Figures 5C, D). Qualitatively, the co-culture clusters remained well-suspended in Z throughout the course of the experiment.

**Figure 5:**
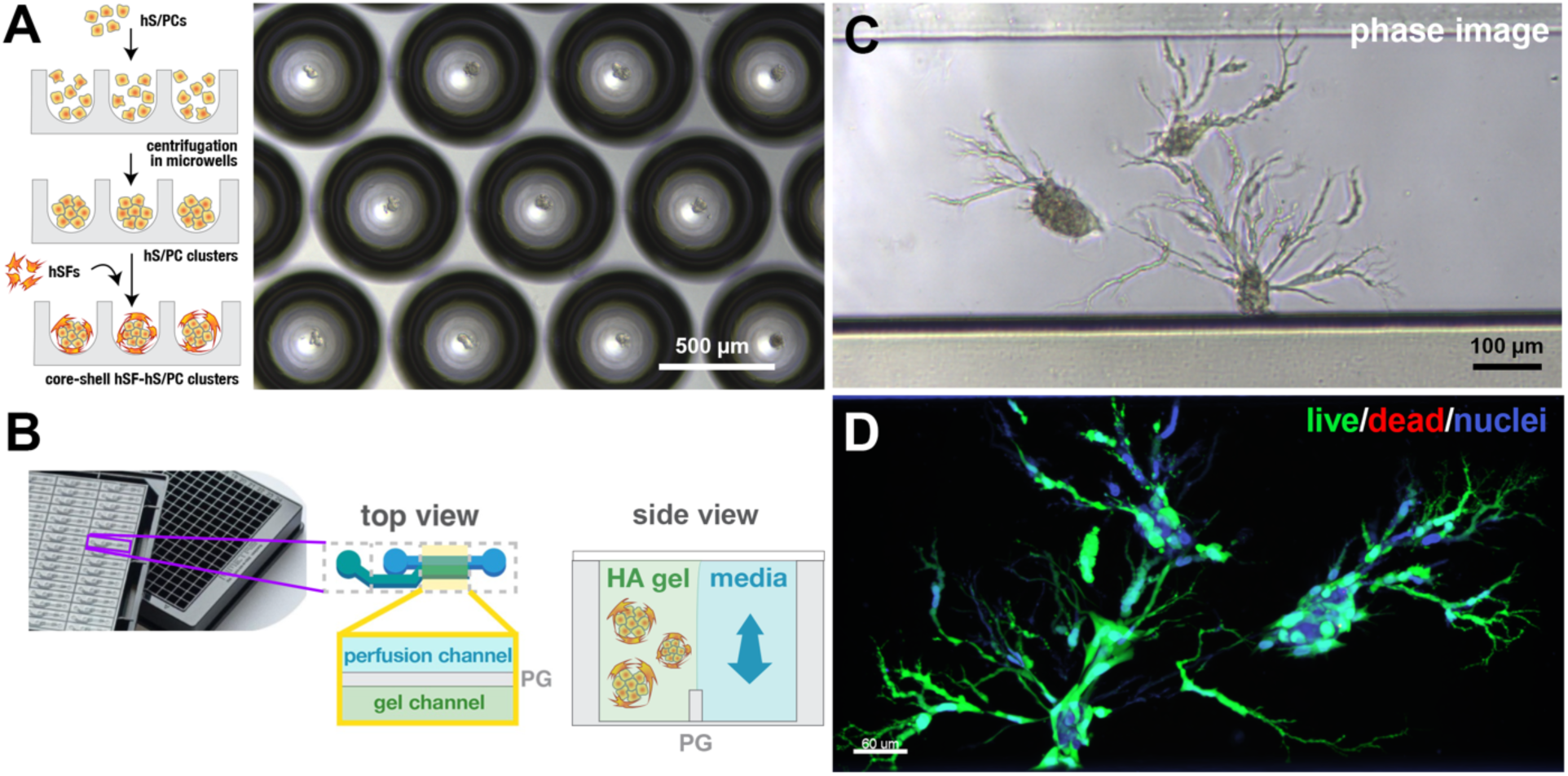
HA hydrogels support human salivary-derived cell co-cultures on microfluidic perfusable high-throughput screening microplates. Core-shell co-cultures of hS/PCs (core) and surrounding hSFs (shell) were pre-clustered, resuspended in hydrogel precursor solutions, and loaded onto a microfluidic plate for extended perfusion culture. Brightfield phase images after 5 days of culture demonstrated a stretched morphology of the peripheral cells, and retention of the internal clusters. Staining for viability and nuclei after 7 days of culture showed high viability (green) with no observed dead cells (red).

## DISCUSSION

From the different hydrogel formulations and compositions that were tested, the following conclusions can be drawn: Gelation kinetics increased with pH with pH 8.0 yielding optimal kinetics; Hydrogel storage modulus, gelation rate, and time to gelation were comparable between PQ and PQ* crosslinked HA-SH gels; Pendant peptide adhesive epitopes Ac-RGD and Ac-YIGSR could also be swapped with comparable rheometric results; PQ:PQ* “safety net” systems yielded a close range of G’ moduli values at varying ratios of high:low MMP degradability crosslinker; PQ* crosslinker exhibited greater resistance to MMP-1 degradation; and HA-SH hydrogel crosslinked with PQ and containing Ac-RGD and Ac-YIGSR epitopes were capable of supporting 3D co-cultures of primary human salivary epithelial cells and salivary fibroblast in a core-shell configuration.

Notably, the pH limits explored in this work for gelation purposes would be prohibitive for extended cell culture. However, we have observed in multiple studies that even fragile primary cells, including the hS/PCs used here, can tolerate a brief excursion to pH 8.0 for gelation purposes, if the full hydrogel construct is returned to a standard buffered media system (pH 7.4) within 60-90 minutes. For the purposes proposed in this work, such as repeat cell dispensing or bioprinting, control over gelation kinetics is critical.

Rheologic data in Figure 2 displayed consistent trends for gelation across multiple compositions. We chose to end data capture at 60 min (3600 s), as our own experience and other past work showed that the majority of thiol-acrylate crosslinking is complete by this time. We observed some variation in the degree of crosslinking completion after 60 minutes, with some compositions reaching a more complete plateau. For the purposes of this investigation, and to avoid potential hydrogel dehydration on the rheometer, we chose to limit the study to the near-maximal level at 60 minutes. Ghosh demonstrated that storage modulus for similar HA-SH/PEGDA systems would increase over an additional 24 hours through disulfide formation, but this variation scaled with the absolute crosslinking density. Within the very soft (<150 Pa) levels of our study, such long-term increases should be <20%.

Similarly, the “safety net” ratios of PQ:PQ* in Figure 3 A,B did not generate perfectly overlapping rheometric data. However, such variability was relatively small, and could have been due to the slight difference in peptide length, the inherent dispersity of the SVA-PEG-acrylate, imperfect coupling or purification of the di-PEGylated peptide crosslinkers, and/or combinations of these effects. We necessarily focused on PQ:PQ* compositions with lower PQ* (0%, 25%, 50%), as these better support HS27a stretching and survival. Modest variation in rheometric data also was observed for exchanges of the pendant Ac-RGD and Ac-YIGSR epitopes, even though they are commercially-produced small molecule compounds.

The MMP degradation data in Figure 3c is an interesting contrast to one of our prior systems (Martinez et al. 2021), which compared similar HA hydrogels, formed with either 100% PQ crosslinker, 100% PQ*, or 100% PEGDA. In that work by Martinez et al, gel pucks *expanded* in size when incubated with MMP-2, while the present work showed that susceptible gel pucks *contracted*. We propose that this contrasting behavior is likely due to the significantly higher PEG content in the Martinez work, in which every pendant RGD was tethered to the network via a PEG oligomer, whereas the present manuscript describes systems with direct N-terminal acrylates for epitope tethering. The Ac-PEG-RGD systems in Martinez exhibited significant network swelling, even prior to enzyme exposure, and we hypothesize that cleaved crosslinkers may liberate PEG chains to occupy more free space, and attract more water molecules; similar expansive swelling behavior was observed by Lutolf for all-PEG hydrogel networks (Lutolf et al. 2003). The Ac-PEG-RGD networks in Martinez contained ∼7-20x more moles of PEG oligomers than the Ac-RGD networks presented here. The contraction of the present networks after MMP-based degradation may be due to residual available thiols along the HA-SH backbone, which may have new freedom to explore space and form new disulfide bonds as covalent crosslinks are cleaved.

The modifications introduced in this work highlight the modularity of the HA hydrogel system, and its tunability to meet the needs of an array of cell lineages for 3D culture applications. When employing such hydrogels in applications like bioprinting, high-throughput screening, or similar needs, predictable material performance is critical. For example, excessive hydrogel swelling (due to excessive PEG or chain cleavage) could lead to occlusion of a microfluidic channel in a lab-on-a-chip device, impacting drug diffusion and outcome measurements. Similarly, premature collapse of the hydrogel matrix on such a device could dramatically change the encapsulated cells’ phenotype, if they sink through a soft hydrogel matrix and settle onto a hard, glass substrate. Customization of an intended environment is a key advantage of such hydrogels, and preservation of that environment can be just as important. The utility of the “safety net” compositions is found in their preservation of both the phenotype and predictable location of encapsulated cells. For example, a matrix with negligible degradability (e.g. 100% PEGDA, or 100% PQ*) yields atypical rounded fibroblasts, limiting cell-matrix interaction and often leading to anoikis. This behavior, which drove other teams to generate similar MMP-degradable matrices (Lutolf et al. 2003; Lei et al. 2011; Kim et al. 2008), anticipated the broad MMP production expected from mesenchymal cells (Zhang et al. 2006). We hypothesized that a pre-programmed intermediate level of matrix degradability would preserve cell behavior, but decrease the rate of cell sinking. In vitro screening applications with imaging rely on predictable cell locations within the region of interest.

Although this work only utilized PQ and PQ* crosslinker molecules, and Ac-RGD and Ac-YIGSR as adhesive epitopes, there are multiple other crosslinker molecules, pendant ligands, or cytokine depots that could be incorporated in future research. For any application seeking to promote more accurate and useful 3D cell culture, work to make the hydrogel more physiologically representative, tailored for cellular response, and reproducible in physical properties will yield great dividends into the future.

## CONCLUSION

In this study, HA-SH hydrogel networks were designed to access a range of moduli, incorporating multiple levels of biorelevant epitopes and crosslinkers. Gelation kinetics were found to increase with increasing pH, and optimized near pH 8.0 to enable fast cell encapsulation. Rheologic measures demonstrated that hydrogel storage modulus, gelation rate, and time to gelation were comparable between PQ and PQ* crosslinked HA-SH gels with high/low MMP degradability. Pendant peptide adhesive epitopes Ac-RGD and Ac-YIGSR could be swapped with comparable rheometric results. By varying ratios of high:low MMP degradability crosslinker, we generated hydrogels with a similar final modulus, but tailored levels of MMP susceptibility. Such “safety net” hydrogels could have value in extending cell encapsulation time in 3D in vitro systems, as we demonstrated for bone marrow-derived fibroblasts on a perfused microfluidic drug screening system. HA-SH hydrogels crosslinked with PQ and containing Ac-RGD and Ac-YIGSR epitopes were capable of supporting 3D co-cultures of human salivary epithelial cell clusters and surrounding salivary fibroblasts. The compositions introduced in this work may yield significant value toward preserving phenotypic cell behavior while retarding hydrogel degradation, in applications such as drug screening, for which kinetic control is critical.

## Author Contributions

Russell T. Wilson: Contributed to conception, design, data acquisition and interpretation, drafted, and critically revised the manuscript. Andrei Bonteanu: Contributed to conception, design, data acquisition and interpretation, drafted, and critically revised the manuscript. Daniel A. Harrington: Contributed to conception, design, data acquisition and interpretation, contributed to statistical analyses, drafted, and critically revised the manuscript. All authors gave their final approval and agree to be accountable for all aspects of the work.

## Acknowledgements

We offer our sincere thanks to individuals who contributed to this work. Dr. J. Nathaniel Holland provided expert support for statistical comparisons. Drs. Josh Valdez and Bowman Bagley at Advanced BioMatrix provided technical support in the HA materials used in this work. Drs. Jeryl D. English and F. Kurtis Kasper critically reviewed the study proposal and provided writing assistance and editing support. The Farach-Carson and Wu laboratories at UTHealth Houston School of Dentistry collectively contributed to data analysis and experimental design. Particular thanks to Caitlynn Barrows for her contribution of isolated hS/PCs, and Dr. Mariane Martinez for her contribution of isolated hSFs. Dr. Kristin Bircsak at MIMETAS US provided expert input in data analysis and experimental design, and access to the ImageXpress imager for data collection in Figure 4.

## Declaration of Conflicting Interests

None.

## Funding

This project was funded by NIH/NCI SBIR Phase II Contract 75N91019C0004 and NIH/NIDCR R01DE032364. UT START funds supported the rheometer, confocal microscopy, and 3D imaging facilities.

## Notes

### Competing Interest Statement

Mimetas US manufactures and sells the Organoplate used in this work, and provided use of an imager for one figure in the manuscript. Although acknowledged for their contribution, no Mimetas employees are authors on this paper, nor did they influence the data, its analysis, or the content of the paper in any way.

### Summary of Updates

Revision on funding sources to include appropriate grants and core facilities.

