## Supplemental Information for "Hyaluronan Hydrogel “Safety Nets” for 3D Cell Culture Applications"

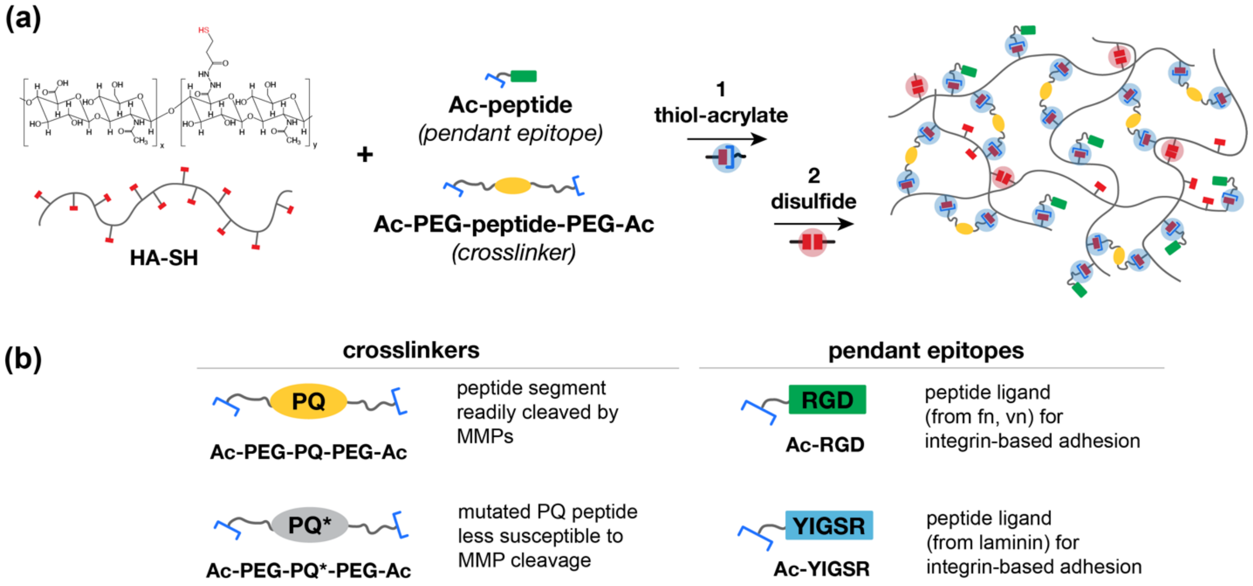


**Appendix Figure 1.** (a) Hyaluronic acid (HA) hydrogels are formed from thiolated derivatives (HA-SH), which create a hydrogel network via a two-step crosslink process: thiols react quickly with acrylates on bifunctional crosslinkers, and remaining thiols progressively form disulfide bridges. Pendant epitopes are added concurrently, and integrate into the backbone through the same process. (b) Acrylated peptidic crosslinkers and pendant epitopes were used in this study to affect network degradability and to enhance integrin-based adhesion by cells. "PQ" and "PQ*" crosslinkers reflect those with high and low susceptibility, respectively, to MMP-based degradation. "RGD" and "YIGSR" pendant groups reflect adhesive ligands present in fibronectin and laminin, respectively, and in other ECM proteins.

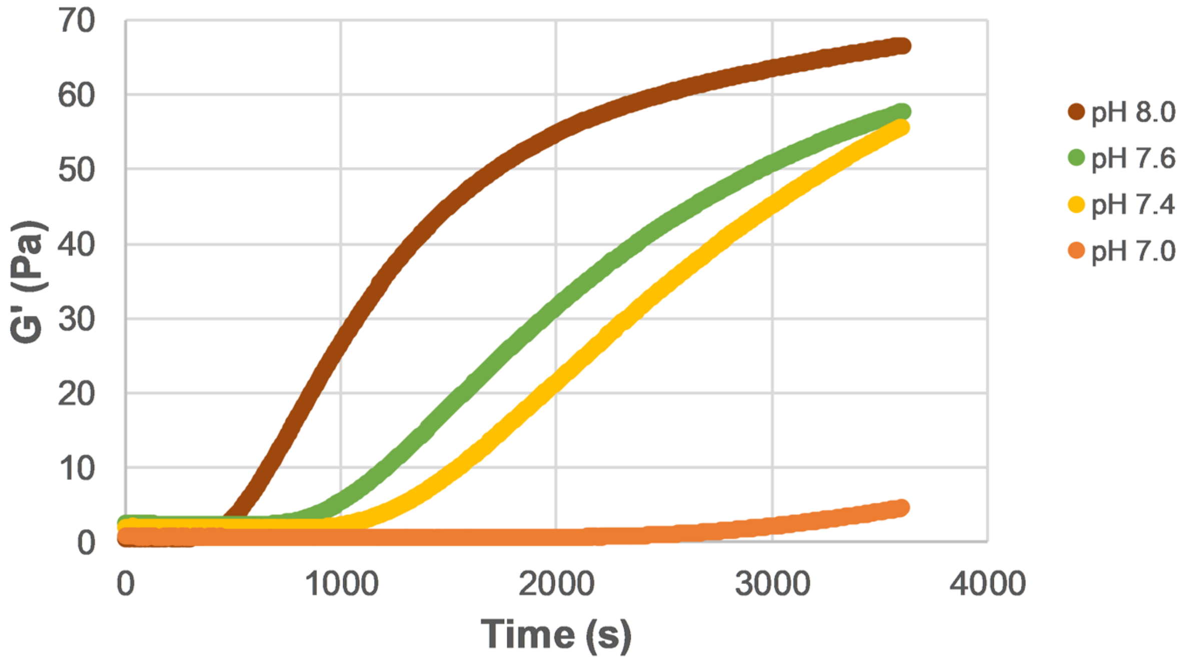


**Appendix Figure 2.** Crosslinking rate for HA-SH hydrogels increases as a function of pH. Rheometric data is shown for HA-SH hydrogels, functionalized with 3.0 mM Ac-RGD and 0.55 mM PQ, at pH ranging from 7 to 8.

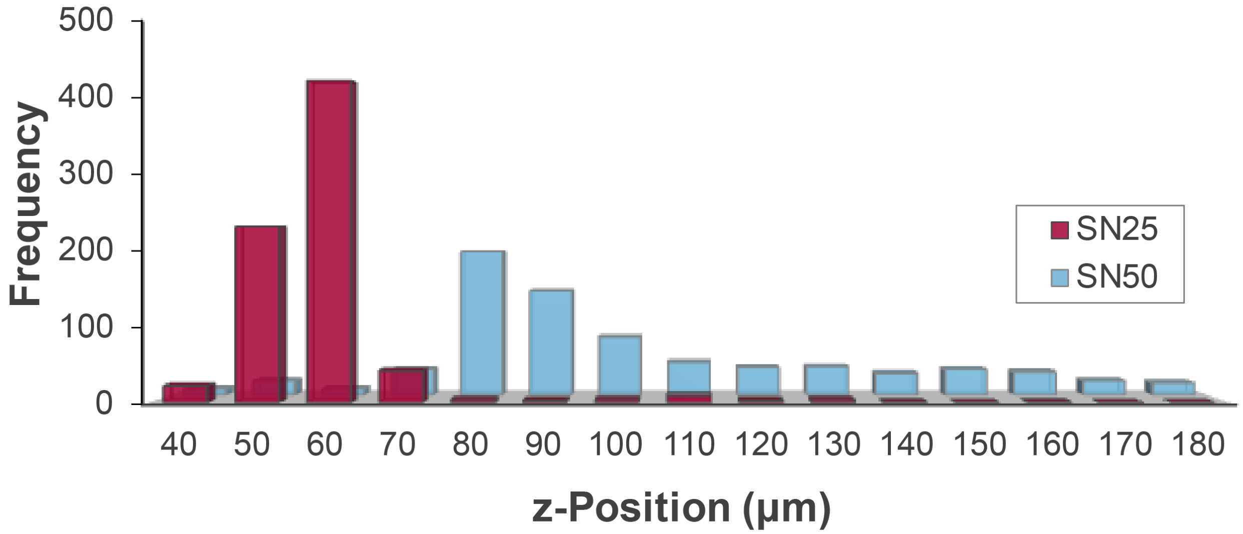


**Appendix Figure 3.** A histogram of HS27a cell location in Z within the GC at day 10 identified that cells within SN25 hydrogel had preferentially fallen to a lower z position, while cells in SN50 retained a more even distribution, with a higher average height within the 200 μm channel.

Appendix Table 1: Tukey's post-hoc comparisons of changes in hydrogel diameter over time, after immersion in MMP solution.

| Comparison | p value | Identifier |
|---|---|---|
| PQ*25 - PQ*0 | 0.51887 | b |
| PQ*75 - PQ*0 | < 0.001 | *** |
| PQ*100 - PQ*0 | < 0.001 | *** |
| PQ*75 - PQ*25 | 0.00185 | ** |
| PQ*100 - PQ*25 | 0.02596 | * |
| PQ*100 - PQ*75 | 0.86482 | a |

### MATERIALS AND METHODS

#### Hydrogel Components

Thiol-modified hyaluronic acid (HA-SH), at average thiolation levels of 35%, or 5 mM of thiols, and degassed water were purchased from Advanced BioMatrix. The following peptides were synthesized commercially by GenScript (>95% purity) with N-terminal acrylation or acetylation, and C-terminal amidation where noted: acryl-GRGDS-amide (Ac-RGD), acryl-YIGSR-amide (Ac-YIGSR), acetyl-KGGGPQGIWGQGK-amide (pre-PQ), and acetyl-KGGGDQGIAGFGK-amide (pre-PQ*). The pre-PQ and pre-PQ* were dissolved separately at 20 mg/mL in 3 mL of 20 mM HEPBS buffer (Santa Cruz Biotech) supplemented with 100 mM NaCl, 2 mM $CaCl_2$, and 2 mM $MgCl_2$. Acryl-PEG-succinimidyl valeric acid (SVA) (Laysan Bio, 3400 Da MW, >95% purity) was then dissolved in 3 mL buffer in stoichiometric molar ratios of acryl-PEG-SVA to peptide of 2.1:1. Both solutions were then titrated to pH 8, combined, and blanketed with argon gas before vortexing for 1 minute. The reaction was placed on a nutating 3D shaker at 4°C and the pH was monitored and re-adjusted to pH 8 every 15 minutes until stable. The reaction was then left to complete overnight. The product was then dialyzed in a 6 kDa – 8 kDa MWCO membrane (Repligen SpectraPor) against 4L of deionized water and agitated with a magnetic stirrer. The dialysate was replaced at least 8 times, each replacement providing at least 2 hours to ensure equilibrium was reached. After dialysis, the product was sterile filtered using a 0.22 µm PES filter and subsequently lyophilized. Final PEGylated crosslinker and binding ligand products were as follows: acryl-PEG-PQ-PEG-acryl (PQ) and acryl-PEG-PQ*-PEG-acryl (PQ*). PEGylation was verified by MALDI-TOF (Bruker AutoFlex II) (PQ: Mn ~8111 g/mol, PQ*: Mn ~8033 g/mol). The lyophilized powder was stored at -20°C blanketed by argon and allowed to warm to room temperature before use.

#### Hydrogel Preparation

Lyophilized HA-SH (10 mg) in amber vials under inert nitrogen blanket was reconstituted through syringe addition of 0.9 – 1.2 mL of degassed water (DGW) to achieve a target active thiolation level of ~35%, or 5 mM of thiols. Contents were vortexed, incubated in a 50°C water bath for 45 minutes, and maintained in a 50°C sonicating water bath for 15 minutes to ensure dissolution, then titrated to pH 8 through addition of 2.5 – 3.5 μL of 1 N NaOH. PQ and/or PQ*, and either Ac-RGD or Ac-YIGSR were dissolved in PBS to make stock solutions at 4.4 mM for PQ/PQ* and 18 mM for Ac-RGD/Ac-YIGSR then titrated to pH 8. These stock solutions of acrylate-bearing moieties were then further aliquoted into "mastermixes" to reach the desired molar combinations of crosslinker and pendant ligands (e.g. final concentration of 0.5 mM crosslinker, using 25% PQ, 75% PQ*, and 3.0 mM Ac-RGD) with further dilution as needed. Prior to loading, each mastermix solution was combined 1:2 (v/v) with HA-SH solution, mixed thoroughly but carefully to avoid bubbles, and loaded onto the rheometer geometry. In pH-specific studies, the above reagents were pH-adjusted using μL increments of 1N HCl or 1N NaOH until reaching the desired pH.

#### Rheometry

Hydrogel variants were characterized mechanically using a TA Instruments Discovery HR-3 Hybrid Rheometer using a 20 mm parallel plate geometry (1 mm gap distance) with an upper solvent trap filled with water; a lower circulating Peltier plate maintained at 37°C; and a surrounding moisture-retaining enclosure. Hydrogel precursor solutions (i.e. reconstituted HA-SH and crosslinker/ligand solution) were titrated to pH 8 (with either 1N NaOH or 1N HCl) unless otherwise noted, mixed evenly, and immediately placed on the geometry for assessment. Measurements were taken at a constant 2% strain and at a recording frequency of 1 Hz. Storage moduli curves reported here are the final values after 60 minutes of measurement during gelation unless otherwise noted.

**Hydrogel Puck Degradation**

Hydrogel precursor solutions were prepared at the desired concentrations and safety net ratios, then combined, mixed evenly, and placed into silicon well molds to produce hydrogel pucks ~6 mm in diameter. After being left to complete gelation over 2 hours in a humidified incubator at 37°C, pucks were carefully removed from the molds, immersed in a PBS solution (Corning phosphate buffered saline without calcium and magnesium) within 12-well multiwell plates, and allowed to equilibrate overnight in a humidified incubator at 37°C. After overnight equilibration, PBS was aspirated carefully from each well, and replaced with a 1 µg/mL solution of human MMP-1 (Sigma-Aldrich SRP3117) in PBS. Samples were returned to the incubator and pucks were imaged daily, using a Nikon SMZ800 microscope at 1x magnification and Nikon Digital Sight DS-Fi2 camera to document changes in hydrogel diameter over 9 days. MMP-1 solution was refreshed in each well at 5 days. Puck diameter was measured digitally using NIS-Elements AR 4.13.04 64-bit software. Puck diameter was normalized to the initial puck diameter for data analysis.

**3D Cell Culture and Imaging – Bone Marrow-Derived Fibroblasts**

HS27A human bone marrow-derived fibroblasts (White, male) were obtained from the American Type Culture Collection (ATCC CRL-2496) and cultured and expanded in RPMI-1640 medium (ATCC 30-2001) supplemented with 10% (v/v) heat inactivated fetal bovine serum (Bio-Techne S11150) and 1% (v/v) penicillin-streptomycin (“complete” RPMI media), on standard tissue culture polystyrene, incubated at 37°C / 5% CO2 / 95% RH. Cells, passages 10 – 20 were split at 80% confluence using 0.25% Trypsin-EDTA and following standard protocols. E2-Crimson fluorescent HS27A were generated via lentiviral transduction using multiplicity of infection (MOI) of 10. Second generation lentivirus was produced following standard protocols and using a transfer plasmid containing E2-Crimson and puromycin resistance (Addgene #139445). Functional titer was measured by transduction of serial dilutions of lentivirus stock. Transduced HS27A were then maintained for 2 weeks under 2 μg/mL puromycin selection to eliminate cells that did not receive the packaged transgene.

For HS27a encapsulation, HA-SH solution (8 – 11 mg/mL in PBS) was prepared by adjusting pH to 8.0, adding 3 mM Ac-RGD in a minimal volume, and incubating the solution overnight. On the following day, HS27a cells were counted, pelleted down, and gently resuspended in the HA-SH solution, at 5E6 cells/mL. Crosslinker in solution was added at a final concentration of crosslinking acrylate groups (2 groups per crosslinker) of 0.5 mM. Two PQ* hydrogel formulas were used at 25:75 and 50:50 molar ratios of PQ*:PQ. The hydrogel solution was gently mixed and loaded into the gel channel of a Mimetas 2-lane OrganoPlate as per manufacturer recommendation. Typical loading volumes were 1.2 μL per channel. OrganoPlates were placed in a humidified incubator for 60 minutes to allow the HA-SH hydrogel to fully solidify within the gel channel. Each 4-well chip was visually inspected under a light microscope and any chips that overflowed or did not fill to completion were noted and excluded from use. 50 µL hS/PC media was then added to the perfusion inlet of each chip. The fully loaded plate was placed on a continually alternating rocker (7 degree, 8 minutes per cycle) in the cell culture incubator to perfuse. Media was exchanged every 3 days by aspirating all media from inlet and outlet wells, taking care not to aspirate the perfusion channel itself. 100 µL of complete media was then added to the inlet well to flush the perfusion channel.

For determining integrity of the gel channel, 100 µL of complete media containing 1 mg/mL of 2 MDa FITC-Dextran (Millipore Sigma, FD2000S) was added to the perfusion channel and allowed to incubate on the rocker for 15 minutes prior to imaging. Confocal imaging was conducted on a Molecular Devices ImageXpress Micro Confocal High-Content Imaging System using Meta Xpress software. Z-series images were obtained for the full height of the channels using either a 10x or 20x LWD objective, and a resolution of 1024 x 1024 pixels. Fluorescence was observed using 488 and 640 nm lasers, and standard FITC (525/50) and far red (685/70) emission cubes, respectively.

Confocal z-stack images were reconstructed in 3D in IMARIS version 9.2 (Oxford Instruments). For gel channel degradation analysis, the gel channel was selected as a region of interest (ROI). Volumetric intrusion of FITC-dextran into the ROI was rendered and calculated as a percentage of the gel channel volume, reported as Volumetric FITC-dextran Intrusion. Thresholding was chosen such that background subtraction eliminated the noise associated with the lowest intensity histogram by pixel intensity value. Background thresholding was consistent across the same day timepoints.

For morphological stretching analysis, the aspect ratio of the cells was computed by first constructing a 3D volume surface in IMARIS labeling individual cells. Then, cell axes were manually labeled and bound to each individual cell surface created to ensure accuracy. Aspect ratio was determined by dividing the longest cell axis by the next longest orthogonal cell axis.

**3D Cell Culture and Imaging – Salivary-Derived Cells**

Human salivary epithelial stem/progenitor cells (hS/PCs) and fibroblasts (hSFs) were isolated from healthy parotid tissue acquired during tumor resection on head and neck cancer patients, using protocols approved by institutional review boards at Rice University, University of Delaware, and University of Texas Health Science Center at Houston (IRB# HSC-DB-16-1060). hS/PC isolation followed previously described methods (Wu et al. 2018). Briefly, parotid tissue was cleaned, minced, and plated onto tissue culture flasks. Cell outgrowth from explants were progressively detached with light trypsinization, and expanded separately. hS/PCs were obtained using a serum-free Williams-E base media, as described (Wu et al. 2018). hSFs were obtained by a similar method, using complete RPMI 1640 medium supplemented with 10% fetal bovine serum, penicillin (100 U/mL), streptomycin (100 µg/mL), and fungizone (1% vol/vol).

hS/PCs were clustered using an ultra-low attachment (ULA) microwell plate (Corning Elplasia 4441), following manufacturer's directions. hS/PCs in 1mL of hS/PC media were added to each well, such that each microwell cavity contained ~34 cells and 1 mL total media. Each well was mixed by gentle pipetting to ensure even dispersion of cells in the media. The cluster plate was then placed in the incubator for 15 minutes to allow the cells to settle into the microwells, after which the clustering plate was incubated on an XY rotator at 60 rpm for 48 hours to allow the clusters to condense. After 48 hours of clustering, hSFs in 0.5 mL hSF media were added to each well such that there were 17 hSF cells per microwell. The cluster plate was then placed in the incubator for 15 minutes to allow the cells to settle into the microwells, after which the clustering plate was incubated on an XY rotator at 60 rpm for an additional 24 hours to allow the fibroblasts to wrap around the epithelial core. After core-shell clustering, the clusters were harvested by gentle pipetting, pelleted, and resuspended in HA-SH hydrogel containing 5 mM thiols, 0.5 mM PQ acrylates, and 2 mM Ac-RGD and 2 mM Ac-YIGSR.

The hydrogel solution with cells was then loaded into a Mimetas 2-lane OrganoPlate, in a similar manner as for HS27a cells, at a target concentration of 15 clusters per imaging window.

After 2 and 7 days of culture, media was aspirated from the inlet and outlet wells of the OrganoPlate channels and replaced with 100 uL of complete media containing 10 μM Hoechst 33342, 1 μM Calcein AM, and 2 μM Ethidium Homodimer III, to fluorescently identify cell nuclei, live cells, and dead cells, respectively. The OrganoPlate was returned to the rocker under incubation for 60 minutes, then removed for imaging. Plates were imaged on a Nikon A1-R confocal microscope using NIS-Elements 5 software. Z-series image stacks were obtained for the full height of the channels using either a 10x or 20x LWD objective, resonance scanning, NIS denoise.ai to remove background noise, a Z-step size of 0.9 μm (20x) or 3 μm (10x), and a resolution of 1024 x 1024 pixels. Fluorescence was observed using 488 and 561 nm lasers, and standard FITC (525/50) and TRITC (600/50) emission cubes, both with GaAsP detectors. 3D image reconstruction was performed in either NIS-Elements software, or Bitplane IMARIS software, as needed.

**Statistics**

Studies involving the physical response of hydrogel pucks to MMP incubation were completed with N=3 pucks per condition. The data on these studies are presented in Figure 4 as the mean of these technical repeats, with error bars representing a 95% confidence interval, assuming a Student's T distribution. 1-way ANOVA, with repeated measures comparison of the specimens, and Tukey's multiple comparisons test, were performed using R statistical software (R Core Team 2020).
